## Supplementary material for "A BRC1-modulated switch in auxin transport accounts for the competition between Arabidopsis axillary buds": Mathematical appendix

Nahas et al., 2025

Starting from the equations used by Prusinkiewicz et al. (2009), the following derivation is performed to obtain the auxin efflux from one of the two buds in a 2-node explant. We assume that the system is symmetrical, so the derivation is the same for the efflux from the second bud, and is only shown for the final equations.

(i) Prusinkiewicz et al. (2009) model the auxin flux between metamers of a plant. Similarly, the model developed here abstracts a plant shoot to the level of axillary buds, but with the extra simplification that we do not explicitly model the section of decapitated main stem between the two buds. We consider only the auxin efflux out of the top and bottom buds, with the stem auxin considered to be equal to the sum of the effluxes from both buds.

(ii) Prusinkiewicz et al. (2009) model canalisation by formulating the change in PIN as dependent on the net auxin flux $\phi$. The net auxin flux from the compartments $i$ to $j$, $\phi_{i\to j}$, is equal to the flux from $i$ to $j$ minus the flux from $j$ to $i$. Each of these unidirectional fluxes depends on the concentration of auxin and PINs, with some diffusion D, which occurs because of apolar auxin transport. From Prusinkiewicz et al. (2009):

$$\frac{d\,[PIN_{i\to j}]}{dt} = \rho_{i\to j}\frac{\Phi^n_{i\to j}}{K^n + \Phi^n_{i\to j}} + \rho_0 - \mu\,[PIN_{i\to j}] \qquad \text{if } \Phi_{i\to j} \geq 0, \tag{1}$$

$$\frac{d\,[PIN_{i\to j}]}{dt} = \rho_0 - \mu\,[PIN_{i\to j}] \qquad \text{if } \Phi_{i\to j} < 0, \tag{2}$$

where $PIN_{i\to j}$ is the surface concentration of PINs transporting auxin from metamer $i$ to $j$, $\rho_{i\to j}$ is the maximum rate of auxin flux-dependent allocation of PINs in the direction $i$ to $j$, $\phi_{i\to j}$ the net flux of auxin from metamer $i$ to $j$, $\rho_0$ is the flux-independent rate of PIN allocation in either direction, and $\mu$ represents the decay or removal of PIN from the metamer surface.

Using net fluxes requires having two conditions for both positive and negative flux from $i$ to $j$. To remove this requirement, we replaced the net flux with a biochemically plausible formulation of the flux ratio between efflux and influx proposed by Cieslak et al. (2015). They obtain this by creating a reaction network where the flux ratio is measured by the concentration of tally molecules X produced each time auxin is transported out of a compartment or, for our purposes, out of the bud. Tally molecules Y are produced each time auxin is transported into the bud. $J_{L\to R}$ is the flux of auxin out of the bud (or efflux), and $J_{R\to L}$ the flux into the bud (or reflux). Y catalyses the turnover of X at rate $v_{XY}$ . Additionally, X and Y are independently degraded at rate $\mu_X$ and $\mu_Y$, respectively. At steady state, the system behaves in the following manner, which enables us to obtain the concentration of X as a proxy for the flux ratio (equations 4.3 from Cieslak et al. (2015)):

$$J_{L\to R} - \mu_x[\mathrm{X}] - v_{xy}[\mathrm{X}][\mathrm{Y}] = 0$$
$$J_{R\to L} - \mu_y[\mathrm{Y}] = 0$$

By solving these equations for [X], we obtain: (3)

$$[\mathrm{X}] = \frac{\mu_y J_{L\to R}}{v_{xy} J_{R\to L} + \mu_x \mu_y}$$

With this formulation for the flux ratio in mind, we can now return to equation 4.1 from Prusinkiewicz et al. (2009), and replace the net flux $\phi$ with the flux ratio X from Cieslak et al. (2015) from equation 4.3:

$$\frac{d\,[PIN_{i\to j}]}{dt} = \rho_{i\to j} \frac{\left(\frac{\mu_y J_{L\to R}}{v_{xy} J_{R\to L} + \mu_x \mu_y}\right)^n}{K^n + \left(\frac{\mu_y J_{L\to R}}{v_{xy} J_{R\to L} + \mu_x \mu_y}\right)^n} + \rho_o - \mu\,[PIN_{i\to j}]$$

Simplifying:

$$\frac{d\,[\mathrm{PIN}_{i\to j}]}{dt} = \rho_{i\to j} \frac{(\mu_y J_{L\to R})^n}{(\mu_y J_{L\to R})^n + K^n\,(v_{xy} J_{R\to L} + \mu_x \mu_y)^n} + \rho_o - \mu\,[PIN_{i\to j}]$$

(iii) We rename the unidirectional fluxes $J_{L\to R}$ and $J_{R\to L}$. $J_{L\to R}$ is renamed as the flux E out of the first bud, and F for the flux out of the second bud. The unidirectional flux $J_{R\to L}$ corresponds to the flux of auxin entering the bud, which is proportional to the auxin in the main stem. We assume that, since our explant consists of only two buds on a decapitated main stem, the influx is proportional to the sum of effluxes from both buds, i.e., $J_{R\to L} = a(E+F)$, where $E+F$ are the sum of effluxes from both buds. The parameter $a$ is a reflux coefficient, determining how well auxin moves from the stem into the bud. Experiments have shown that radiolabelled auxin applied to the stump of a decapitated stem travels down the stem without accumulating in the bud (Brown et al., 1979; Everat-Bourbouloux and Bonnemain, 1980; Booker et al., 2003), so in practice it would be negligible. This simplification enables us to represent the source-sink relationship between the bud and the stem without explicitely representing the stem. The parameter $a$ can be conceptualised as influencing the strength of the source-sink relationship. We also assume that, despite auxin being transported basipetally, auxin transport canalisation enables both the top and the bottom bud to inhibit each other equally. This assumption is justified given the closeness of the two axillary buds on two-node explants. In addition, *in planta* some of the inhibition from the bottom bud to the top bud may be acting via the acropetal movement of strigolactone, though for simplicity, we model 2-node explants as symmetrical.

We get:

$$\frac{d\,[PIN_{i\to j}]}{dt} = \rho_{i\to j} \frac{(\mu_y E)^n}{(\mu_Y E)^n + K^n\,(v_{XY} a(E+F) + \mu_X \mu_Y)^n} + \rho_o - \mu\,[PIN_{i\to j}] \tag{4}$$

(iv) The equation above is partly formulated in terms of PIN concentration and partly in terms of auxin efflux. To reformulate it only in terms of efflux, without PIN concentrations, we assume that the concentration of auxin in the bud is constant. This can be justified from evidence of auxin homeostatic feedback (Ljung et al., 2001). The assumption of constant auxin concentration enables the following steps:

The efflux $E$ of auxin out of the bud corresponds to:

$$E = [auxin]([PIN]_{i\to j} + D),$$

where $[auxin]$ is the constant auxin concentration.

Reordering:

$$[PIN_{i\to j}] = \frac{E}{[auxin]} - D$$

Given the constant auxin concentration, the time derivative of $[PIN_{i\to j}]$ is the time derivative of E divided by the bud auxin concentration:

$$\frac{d[PIN_{i\to j}]}{dt} = \frac{\frac{dE}{dt}}{[auxin]}$$

We use this formulation to replace $PIN_{i\to j}$ and $d[PIN_{i\to j}]$ in equation 4.4.

$$\frac{\frac{dE}{dt}}{[auxin]} = \rho_{i\to j}\frac{(\mu_y E)^n}{(\mu_Y E)^n + K^n\left(v_{xY}a(E+F) + \mu_x\mu_Y\right)^n} + \rho_o - \mu\left(\frac{E}{[auxin]} - D\right)$$

$$\frac{\frac{dE}{dt}}{[auxin]} = \rho_{i\to j}\frac{(\mu_y E)^n}{(\mu_Y E)^n + K^n\left(v_{XY}a(E+F) + \mu_X\mu_Y\right)^n} + \rho_o - \mu\frac{E}{[auxin]} + \mu D$$

We now multiply each side by the constant auxin concentration:

$$[auxin]\frac{\frac{dE}{dt}}{[auxin]} = [auxin]\left(\rho_{i\to j}\frac{(\mu_y E)^n}{(\mu_Y E)^n + K^n\left(v_{XY}a(E+F) + \mu_X\mu_Y\right)^n} + \rho_o - \mu\frac{E}{[auxin]} + \mu D\right)$$

$$\frac{dE}{dt} = [auxin]\rho_{i\to j}\frac{(\mu_y E)^n}{(\mu_Y E)^n + K^n\left(v_{XY}a(E+F) + \mu_X\mu_Y\right)^n} + \rho_o[auxin] - \mu E + \mu D[auxin]$$

Let $v0 = \rho_0[auxin] + \mu D[auxin] = (\rho 0 + \mu D)[auxin]$, such that $v0$ is now the basal rate of efflux from the bud.

Let $v = \rho_{i\to j}[auxin]$, such that $v$ is now the maximum rate of efflux of the Hill coefficient.

$$\frac{dE}{dt} = v0 + v\frac{(\mu_y E)^n}{(\mu_Y E)^n + K^n\left(v_{XY}a(E+F) + \mu_X\mu_Y\right)^n} - \mu E$$

(v) We rename the parameters $\mu_X$, $\mu_Y$, $v_{XY}$ from the measure of relative flux to give them a more intuitive meaning:

- $\mu_Y$ is the degradation of tally molecules Y that measure reflux. As $\mu_Y$ increases, the degradation of X decreases, such that the measure of efflux increases. Therefore, we can reformulate $\mu_Y$ as being conceptually the strength of the positive feedback of efflux on itself.

- $\mu_X \mu_Y$ can be grouped into a single parameter C. The parameter space ($\mu_X$, $\mu_Y$) is remapped to the parameter space (S, C), where S $=\mu_Y$ and C $= \mu_X \mu_Y$.

$$\frac{dE}{dt} = v0 + v\frac{(SE)^n}{(SE)^n + K^n\left(v_{XY}a(E+F)+C\right)^n} - \mu E$$

Moving $K$ into the bracket:

$$\frac{dE}{dt} = v0 + v\frac{(SE)^n}{(SE)^n + \left(Kv_{XY}a(E+F)+KC\right)^n} - \mu E$$

(iv) We rescale parameters to simplify the model:

First rescaling: let $K = \frac{K'}{v_{XY}a}$ and $C = C'v_{XY}a$

$$\left(Kv_{XY}a(E+F)+KC\right)^n =$$
$$\left(\frac{K'}{v_{XY}a}v_{XY}a(E+F) + \frac{K'}{v_{XY}a}C'v_{XY}a\right)^n = \quad \left(K'(E+F)+K'C'\right)^n$$

Second rescaling: let $C' = \frac{C''}{K'}$

$$\left(K'(E+F)+K'\frac{C''}{K'}\right)^n = \left(K'(E+F)+C''\right)^n$$

Finally we get:

$$\frac{dE}{dt} = v0 + v\frac{(SE)^n}{(SE)^n + \left(K'(E+F)+C''\right)^n} - \mu E$$

We rename $D = K'$ and $K = C''$ to obtain the final formulation of the model, representing the efflux E out of one bud inhibited by the efflux F from the other bud. We can now also make a second equation for the efflux F out of the second bud, inhibited by efflux E from the first bud:

$$\begin{aligned}\frac{dE}{dt} &= v0 + v\frac{(SE)^n}{(SE)^n + (D(E+F)+K)^n} - \mu E\\ \frac{dF}{dt} &= v0 + v\frac{(SF)^n}{(SF)^n + (D(E+F)+K)^n} - \mu F,\end{aligned} \tag{5}$$

The change in auxin efflux is described by: (i) $v0$, a basal rate of efflux, which corresponds to non-polar auxin transport and/or diffusion of auxin out of the bud, (ii) a Hill function that represents the positive feedback of auxin efflux on itself, mirroring previous models of canalisation, with $v$ as the maximum rate of efflux for the Hill function, $S$ the efficiency of feedback-driven auxin efflux, $n$ the Hill exponent that influences the degree of non-linearity in the feedback, and $K$ a threshold parameter, and (iii) $\mu$, a linear decrease in efflux, which relates to both the removal of PINs from the plasma membrane and the degradation of auxin. The auxin efflux from one bud, conceptually contributing to the auxin concentration in the stem, dampens its own auxin efflux and that of the other bud at a strength that is proportional to $D$ and proportional to the sink strength of the main stem, which is determined by the sum of auxin efflux E and F from each bud. This captures the source/sink dynamics between the buds and the main stem. The modifications performed to develop this simplified model preserve the sigmoidal relationship between auxin flux and the rate of change, which is key to the results obtained by Prusinkiewicz et al. (2009).

To model the growth of buds on 2-node explants, we relate the change in auxin efflux to the growth rate of buds. The relationship between the steady state of auxin efflux E and F for the top and bottom bud, and their growth rate N and M is represented as a Hill function:

$$\begin{aligned} \frac{d[N]}{dt} &= \frac{E^m}{Q^m + E^m} \\ \frac{d[M]}{dt} &= \frac{F^m}{Q^m + F^m} \end{aligned} \tag{6}$$

Where $Q$ and $m$ are Hill coefficients that influence the degree of non-linearity between auxin efflux and growth rate, and $u$ is the maximum of the Hill function. The full model consists of equations 5 and 6, which form two coupled sets of differential equations.
